# Single-cell transcriptome reveals the redifferentiation trajectories of the early stage of *de novo* shoot regeneration in *Arabidopsis thaliana*

**DOI:** 10.1101/2022.01.01.474510

**Authors:** Guangyu Liu, Jie Li, Ji-Ming Li, Zhiyong Chen, Peisi Yuan, Ruiying Chen, Ruilian Yin, Zhiting Liao, Xinyue Li, Ying Gu, Hai-Xi Sun, Keke Xia

**Author notes:** Corresponding authors: Keke Xia, Hai-Xi Sun, Ying Gu. These authors contributed equally to this work (Guangyu Liu, Jie Li, Ji-Ming Li). **Author list:** Guangyu Liu, Jie Li, Ji-Ming Li, Zhiyong Chen, Peisi Yuan, Ruiying Chen, Ruilian Yin, Zhiting Liao, Xinyue Li, Ying Gu, Hai-Xi Sun, Keke Xia.

## Abstract

*De novo* shoot regeneration from a callus plays a crucial role in both plant biotechnology and the fundamental research of plant cell totipotency. Recent studies have revealed many regulatory factors involved in this developmental process. However. our knowledge of the cell heterogeneity and cell fate transition during *de novo* shoot regeneration is still limited. Here, we performed time-series single-cell transcriptome experiments to reveal the cell heterogeneity and redifferentiation trajectories during the early stage of *de novo* shoot regeneration. Based on the single-cell transcriptome data of 35,669 cells at five-time points, we successfully determined seven major cell populations in this developmental process and reconstructed the redifferentiation trajectories. We found that all cell populations resembled root identities and undergone gradual cell-fate transitions. In detail, the totipotent callus cells differentiated into pluripotent QC-like cells and then gradually developed into less differentiated cells that have multiple root-like cell identities, such as pericycle-like cells. According to the reconstructed redifferentiation trajectories, we discovered that the canonical regeneration-related genes were dynamically expressed at certain stages of the redifferentiation process. Moreover, we also explored potential transcription factors and regulatory networks that might be involved in this process. The transcription factors detected at the initial stage, QC-like cells, and the end stage provided a valuable resource for future functional verifications. Overall, this dataset offers a unique glimpse into the early stages of *de novo* shoot regeneration, providing a foundation for a comprehensive analysis of the mechanism of *de novo* shoot regeneration.

## Introduction

In contrast to animals, plants have a remarkable regeneration capability, ensuring developmental plasticity in various environmental conditions. Until now, two modes of regeneration have been described, including indirect *de novo* shoot regeneration and direct shoot organogenesis^1^. Indirect shoot regeneration includes two steps: pluripotent callus induction on an auxin-rich callus inducing medium (CIM) and shoot regeneration on a cytokinin-rich shoot inducing medium (SIM)^1^. Based on plant regeneration ability, *in vitro* tissue culture techniques have been widely used in agriculture^2^. Although many studies have been carried out on plant regeneration, the mechanistic details of shoot regeneration are largely unknown. Shoot regeneration is still a major hurdle in tissue culture^3,4^. Therefore, a comprehensive and deeper understanding of the plant regeneration process is urgently needed.

During *de novo* shoot regeneration, cells undergo complex cell fate transition and reprogramming, and four phases have been defined recently, including pluripotency acquisition (phase I), shoot pro-meristem formation (phase II), the establishment of the confined shoot progenitor (phase III), and shoot outgrowth (phase IV)^5^. These phases involve highly dynamic changes in the transcriptome and reprogramming of cell fate. Recently, several genes and regulatory networks involved in regulating shoot regeneration have been identified. For example, *WUSCHEL* (*WUS*), a core transcriptional factor required for shoot regeneration, is regulated by the dynamic expression of Type-B *ARABIDOPSIS RESPONSE REGULATOR*s (*ARR*s), including *ARR1, ARR2, ARR10*, and *ARR12*^6,7^. The *WOX11–LBD16* pathway, *PLT3, 5, 7-PLT1, 2* pathway, and *WOX5-PLT1, 2-TAA1* pathway promote pluripotency acquisition in callus cells^8-10^, and *PLT3, 5, 7*-*CUC2* pathway promotes *de novo* shoot formation^9^. What’s more, *CUP-SHAPED COTYLEDON 1* (*CUC1*) and *CUC2* are controlled by *ENHANCER OF SHOOT REGENERATION 1* (*ESR1*) for the establishment of shoot pro-meristem^11-13^. However, the knowledge on the regulatory network of *de novo* shoot regeneration is still limited.

In addition, the interaction of multiple cell types and the spatiotemporal dynamic expression of genes also play important roles in controlling *de novo* shoot regeneration^7,14,15^ such as the mutually exclusive and spatiotemporal specific distribution pattern of auxin and cytokinin response cells^14^. Auxin and cytokinin are uniformly distributed at the edge region of the callus cultured on CIM. After transferring callus to SIM, auxin response cells progressively translocate and form the ‘auxin ring’, whereas the cytokinin response cells are restricted to the center of the auxin ring, overlapping with the region of *WUS* expression. These findings suggest that a variety of cell types with unknown characteristics appear progressively during *de novo* shoot regeneration.

Despite extensive studies, many questions related to *de novo* shoot regeneration remain to be answered. For example, to what extent does heterogeneity present in a callus? Which cell type in a callus can regenerate new shoots? How do cell populations change during *de novo* shoot regeneration? What are the expression patterns of genes involved in shoot regeneration? These unsolved questions are largely due to technical limitations. For instance, bulk RNA-seq has been widely used in characterizing molecular changes during *de novo* shoot regeneration, but information related to callus heterogeneity is lost^15,16^. Therefore, to answer these questions mentioned above, new techniques with higher resolution are needed.

Recent advances in single-cell RNA sequencing (scRNA-seq) technology provide unprecedented opportunities for systematically identifying cellular and molecular differentiation trajectories of plant regeneration at the single-cell level^17,18^. The scRNA-seq technique has been widely used in plant development studies, facilitating reconstructing the development trajectories and establishing gene expression regulatory networks of root epidermis^19,20^, lateral root development^21-28^, shoot apex^29,30^, leaf phloem cells^31,32^, guard cells^33,34^, seedlings^35^ and callus^10^, demonstrating its advantages in discovering regulatory elements^36^. Up to now, scRNA-seq has not been used to analyze the complex shoot regeneration process in *Arabidopsis*.

Here, by performing time-series single-cell transcriptome experiments, we revealed the early redifferentiation process of *de novo* shoot regeneration at the single-cell level. First, we successfully reconstructed the early redifferentiation trajectories of *de novo* shoot regeneration. Next, we carried out the cell identities analysis and found seven major cell populations that resembled QC cells and other root cell types. Moreover, we discovered the dynamic expression patterns of canonical regeneration-related genes and also explored potential transcription factor regulatory networks involved in *de novo* shoot regeneration, which provided valuable resources for a deeper understanding of the mechanism of shoot organogenesis.

## Results

### Constructing the early redifferentiation trajectories during *de novo* shoot regeneration from callus cells

During the early period of *de novo* shoot regeneration, we could observe distinct morphological changes. For example, the callus gradually turned compact and green after the callus was transferred to SIM for around 10 days (Supplementary Fig. 1a), indicating that the cell identities had already been reprogrammed. To comprehensively understand cell fate transition during the early stage of *de novo* shoot regeneration, we performed scRNA-seq using root-derived callus cultured on SIM for 0, 2, 4, 6, 10 day(s), respectively (Fig. 1a). For each time point, three replicates were included. After data filtering, we obtained a total of 35,669 cells from the five time points. The cell population from each time point was designated as SIM0, SIM2, SIM4, SIM6, and SIM10, which had 7,571, 9,755, 4,893, 7,795, and 5,655 cells, respectively. In total, 21,129 genes were detected with an average of 1,652 genes per cell.

**Fig. 1.**
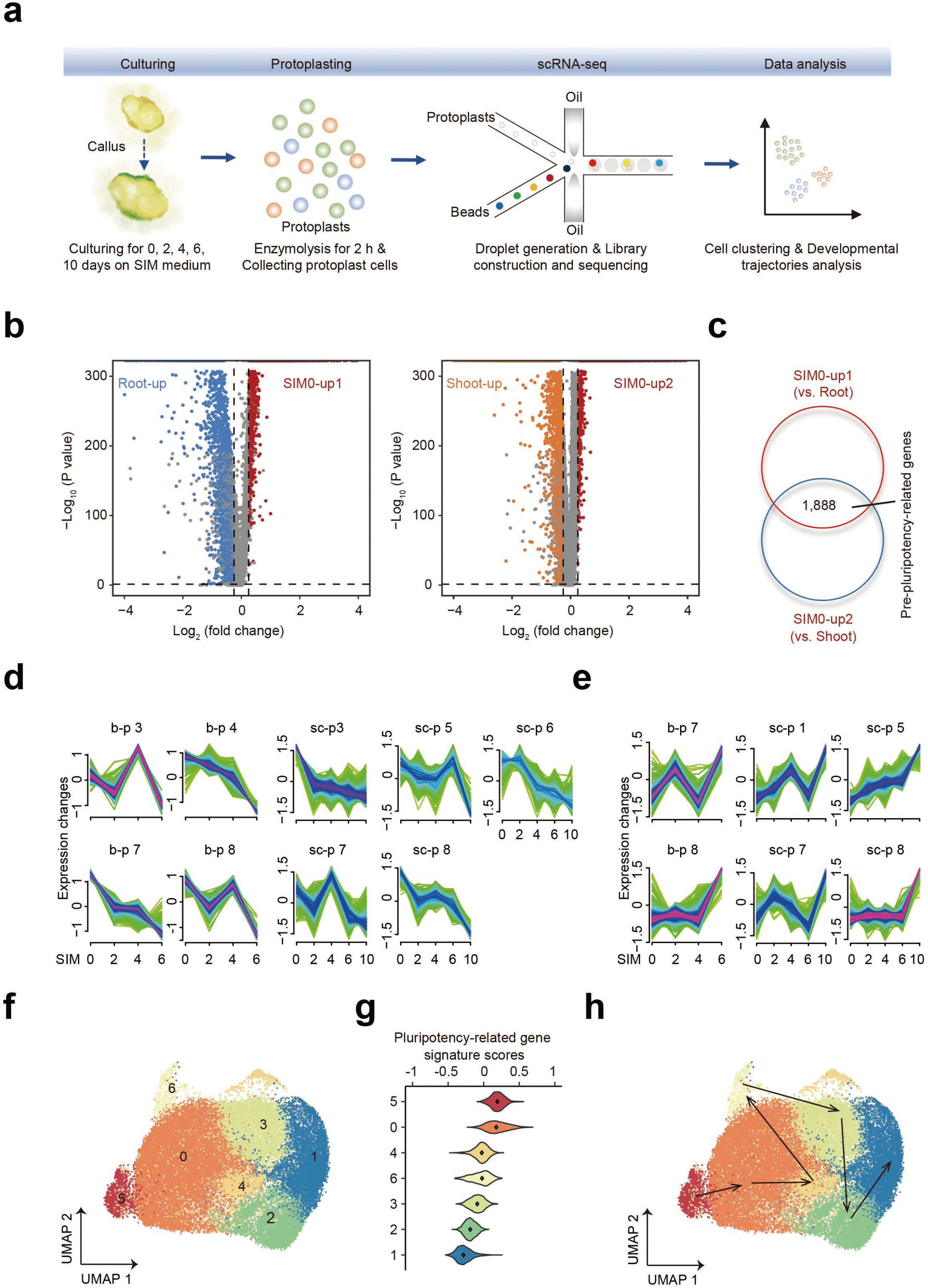
The developmental trajectory during the early stage of *de novo* shoot regeneration. **a**, Schematic representation of the scRNA-seq procedure. **b**, Volcano plots showing the differentially expressed genes (DEGs) in the root (left) and shoot (right) compared with SIM0. The red dots: the up-regulated genes in Day 0, the yellow dots: the up-regulated genes in the shoot, and the blue dots: the up-regulated genes in the root. The gray dots: genes with no significant difference. **c**, The overlap of the up-regulated DEGs in SIM0 compared with the root and shoot. **d**, The pre-pluripotency-related genes with decreased expression patterns in our bulk RNA-seq and scRNA-seq data of samples in Day 0, 2, 4, 6, and 10. **e**, The pre-differentiation-related genes with increased expression patterns in our bulk RNA-seq and scRNA-seq data of samples in Day 0, 2, 4, 6, and 10. **f**, Clustering results of 35,669 cells. **g**, Signature scores of pluripotency-related genes at the single-cell level. **h**, The development trajectory of cell clusters. SIM0 represents the cell population on Day 0. Root-up, shoot-up, and SIM0-up represent up-regulated genes in the root, shoot, and SIM0, respectively. B-p and Sc-p represent patterns in bulk RNA-seq and scRNA-seq, respectively.

To check the reproducibility of the data, we performed correlation analysis among all samples. We found that samples from the same time point were clustered together and showed high similarity (Supplementary Fig. 1b), indicating reproducibility of the data. In addition, to assess data consistency at the gene level among cell populations from different time points, we checked the gene number, the unique molecular identifier (UMI) number, and the proportion of mitochondrial and chloroplast genes. We observed similar numbers for different time points for each of the tested parameters (Supplementary Fig. 1c). Lastly, a comparison of gene expression between scRNA-seq and bulk RNA-seq at day 0, day 2, day 4, and day 6, respectively, also showed the reliability of our dataset (Supplementary Fig. 1d). Taken together, we obtained reliable and reproducible data which were used for further analysis.

As depicted in Supplementary Fig. 1b, the correlation of cell populations changed gradually from SIM0 to SIM10, demonstrating that the gene expression patterns of these cells had changed during *de novo* shoot regeneration. To reconstruct the redifferentiation trajectories during *de novo* shoot regeneration, we first used well-known tools, including monocle 2 and monocle 3^37^. However, we found that the resulting pseudo-time trajectories contradicted with timeline (Supplementary Fig. 2a-e). Thus, monocle 2 and monocle 3 might be not suitable for this dataset. In this study, the root-derived pluripotent callus would eventually develop into the more differentiated shoot apex and shoot tissues. Therefore, we decided to manually select pluripotency- and differentiation-related genes for re-constructing the developmental trajectories.

First, we compared SIM0 scRNA-seq data with the previously reported scRNA-seq data obtained from root^38^ and shoot apex^29^, respectively. We found that there were 2,623 and 2,632 genes up-regulated in callus compared with root and shoot apex, respectively. Then the 1,888 overlapped genes of these two up-regulated genes lists were obtained and designated as pre-pluripotency-related genes (Fig. 1b,c and Supplementary Table 1). Then, we further selected genes that might be involved in pluripotency or differentiation processes by assessing the expression patterns of these 1,888 genes in both bulk RNA-seq datasets and the pseudo-bulk RNA-seq dataset. Among them, 721 genes that had decreasing expression patterns in both the bulk RNA-seq dataset and the pseudo-bulk RNA-seq dataset were found and defined as pluripotency-related genes (Fig. 1d and Supplementary Fig. 3c). Similarly, we obtained 287 genes that had increased expression patterns in both the bulk RNA-seq dataset and the pseudo-bulk RNA-seq dataset and designated these genes as differentiation-related genes (Fig. 1e and Supplementary Fig. 3c). In addition, we also included 61 (6 genes were repetitive with selected pluripotency- and differentiation-related genes) previously reported genes that are involved in shoot regeneration as regeneration-related genes (Supplementary Fig. 3d,e). Finally, a total of 1,063 genes, including 721 pluripotency-related genes, 287 differentiation-related genes, and 55 regeneration-related genes, were selected as input for cell clustering using the Seurat package (Supplementary Table 1). As shown in Fig. 1f, the 35,669 cells were clustered into 7 clusters. Excitingly, expression of pluripotency-related genes in different clusters showed a gradual decrease in the order of cell population 5, 0, 4, 6, 3, 2, and 1 (Fig. 1g), and the order of cell populations largely corresponded to the main populations in each time point (see below), Thus, we reconstructed the redifferentiation trajectories according to cell pluripotency and differentiation, and we defined the redifferentiation trajectories as the following: the cell population 5 were highly pluripotent, which gradually differentiated into cell population 0, 4, 6, 3, 2 and finally developed into more differentiated cell population 1 during the early stage of *de novo* shoot regeneration (Fig. 1h). So far, the scRNA-seq technology facilitated the construction of the landscape of redifferentiation trajectories, which could offer insights into the characteristics of the cell populations during *de novo* shoot regeneration.

### Cell populations resemble QC cells and other known root cell types during the early stage of *de novo* shoot regeneration

Recently, Shin et al. has reconstituted the shoot regeneration process with four phases, including pluripotency acquisition (phase I), shoot pro-meristem formation (phase II), establishment of the confined shoot progenitor (phase III), and shoot outgrowth (phase IV)^5^. Here, to determine the developmental stage of shoot regeneration in our dataset, we assessed the expression pattern of phase-specific genes at the five time points. As expected, phase I-specific genes were expressed, including *WOX5, SCR*, and *PLT1*/*2* (Fig. 2a and Supplementary Fig. 4a). In addition, we detected the expression of phase II-specific genes, such as *PIN1, CUC2*, and *ESR1* (Fig. 2a and Supplementary Fig. 4a). However, we did not detect the expression of phase III-specific genes, such as *WUS* and *CLV3*, and these genes are key shoot meristem marker genes. Moreover, the expression of *WUS* was also not detected in our bulk RNA-seq data (Data not shown). These results demonstrated that the shoot organogenesis process in this study preceded phase III.

**Fig. 2.**
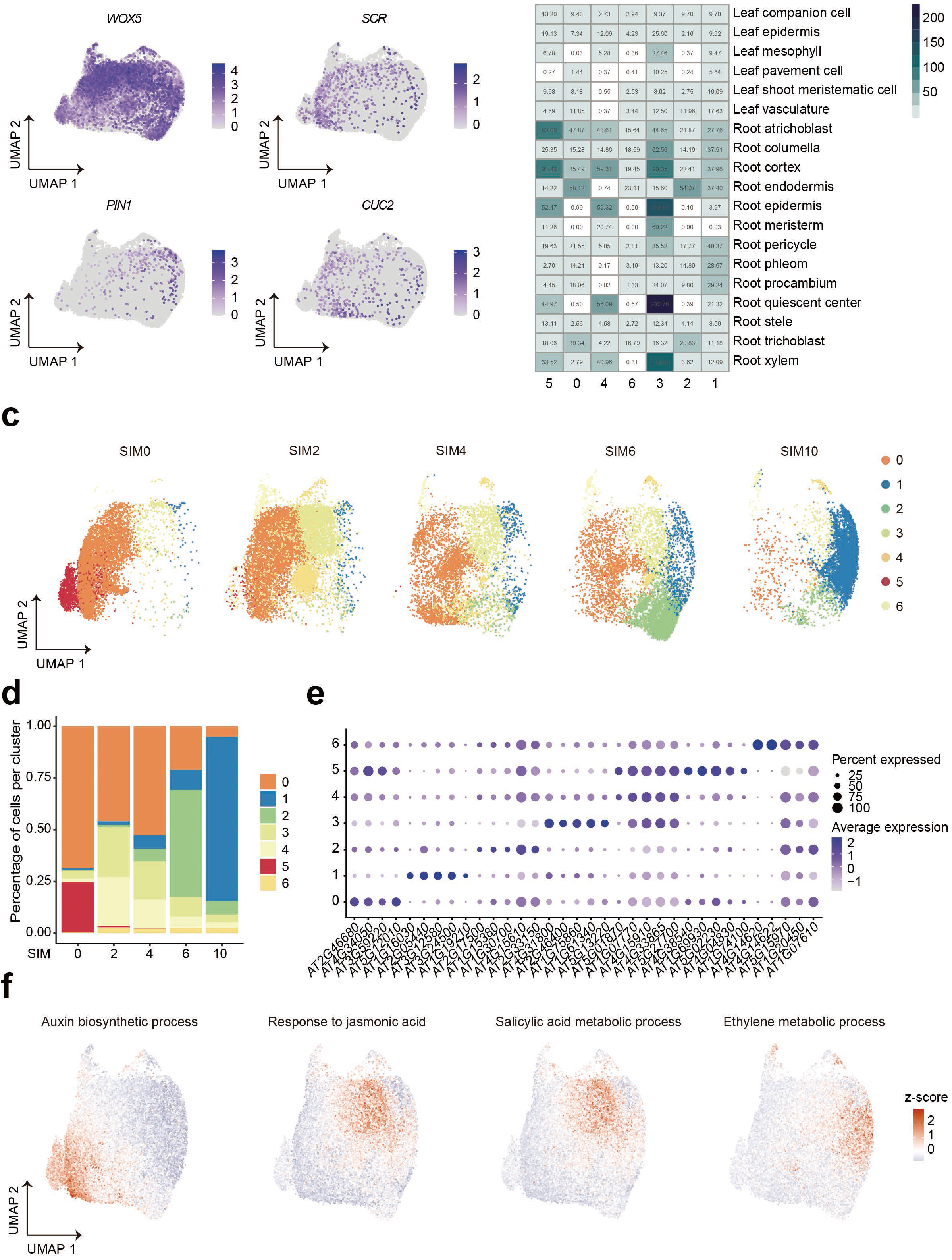
Cell type identification and characterization. **a**, Expression of phase-specific genes at single-cell level during *de novo* shoot regeneration, including *WOX5, SCR, PIN1*, and *CUC2*. **b**, Hypergeometric Distribution Enrichment Analysis (HDEA) of cell populations. **c**, Profiles of cell populations in Day 0, 2, 4, 6, and 10. **d**, The proportion of cluster-based cells in Day 0, 2, 4, 6, and 10. **e**, The expression patterns of the top 5 overlapped genes for each cluster between cluster-specific markers and the reference cell type-specific marker genes. Percent Expressed means percent of gene-expressed cell number in all cells of specific clusters. The average expression means the average expression of genes at the single-cell level in specific clusters. One overlapped marker gene was detected. **f**, Hormone-related GO terms in specific cell clusters, including auxin biosynthetic process in clusters 0 and 5, response to jasmonic acid, and salicylic acid metabolic process in cluster 3 and ethylene metabolic process in cluster 1.

Next, we explored whether the identified populations showed any quantitative affinities to the known cell types. It has been shown that callus has root identity^39^, and root identity marker genes, such as *WOX5, SCR*, and *PLT1*/*2*, were also expressed in our dataset. As *de novo* shoot organogenesis eventually leads to the formation of the shoot, we systematically explored known cell identities, including both root and shoot cell types (Supplementary Table 2). It turned out that cell populations in our dataset were more similar to root cell types rather than leaf cell types (Fig. 2b). For example, cell populations 0, 2, and 6 had the highest endodermis scores, while cell populations 4 and 5 resembled root epidermis and cortex, respectively. Surprisingly, cell population 3 presented the highest score for quiescent center (QC) cells. Consistently, QC-like cells have also been found in hypocotyl-derived callus in a recent study^10^. Lastly, cell populations changed dramatically at day 10, with the majority of cells belonging to cell population 1 (Fig. 2c,d). We found that cell population 1 had an affinity to pericycle identity (Fig. 2b). It has been shown that the pericycle cells have the organogenetic capacity and are the source of lateral roots^40^, suggesting that the pericycle identity of population 1 might play an important role in the shoot regeneration process. In addition to pericycle cells, cell population 1 also showed high scores for other root cell types, such as the cortex, columella, and endodermis cells, suggesting that the composition of cell population 1 might be complicated.

To further validate cell population identities, we determined the overlap between cluster-specific markers in our dataset and the reference cell type-specific marker genes (Supplemental Table 2). And then we generated the heatmap using the top 5 overlapped marker genes. As shown in Fig. 2e, the top 5 overlapped genes for cell clusters 5, 3, and 1 had relatively high specific expression patterns. We also observed that the marker genes of cell cluster 0 were also highly expressed in cell population 5, suggesting that these two populations were similar. In addition, we found *AT4G14620* and *AT4G14622* were specifically and highly expressed in cluster 6. In conclusion, these specifically expressed marker genes not only confirmed the reliability of the cell type identification but also provided potential marker genes and candidate genes for future functional studies, such as *AT4G14620* and *AT4G14622*.

Then we investigated gene ontology (GO) enrichment in the top 100 gene markers (Supplemental Table 2) for each population and found specific features (Supplementary Fig. 4b and Supplemental Table 2). Interestingly, multiple plant hormone pathways were specifically enriched in certain cell populations (Supplementary Fig. 4b). It is well-known that phytohormones, such as auxin, JA, SA, and ethylene, play important roles during plant regeneration^5,41^, when and where these phytohormones play their roles remain unclear. We found that the GO terms associated with populations 5 and 0 were highly similar, and the auxin biosynthesis process was specifically enriched in these cell populations. For cell population 3 (QC-like cells), biological processes related to jasmonic acid (JA) and salicylic acid (SA) were specifically enriched (Fig. 2f). In addition, the GO terms related to the ethylene metabolic process were enriched in cell population 1 (Fig. 2f). Here, based on the re-constructed trajectories using scRNA-seq, we were able to demonstrate the spatiotemporal roles of these phytohormones during shoot regeneration.

In addition to the enrichment of phytohormone-related biological processes, other GO terms were also enriched in specific cell populations. For example, the amine metabolism process in cell populations 5 and 0 (Supplementary Fig. 4c), suggesting that the amine metabolism process might be involved in regulating the formation of callus. Interestingly, we also observed stress-related process was specifically enriched in cell population 1 (Supplementary Fig. 4b). Recently, Zeng et. al found that the stress-related signal was overrepresented in the shoot stem cell niche, and it was necessary to maintain *Arabidopsis* shoot stem cells^42^. Here, the enriched stress response processes in cell population 1 appeared before the formation of the shoot apex, indicating the stress signaling might play a role in regulating the formation of shoot stem cells.

### Dynamic expression patterns of regeneration-related genes along the redifferentiation trajectory

So far, many key regulatory genes have been discovered during the shoot regeneration process^5,43^, but few studies have focused on the dynamic expression patterns of these genes. In this study, we detected the expression of 75 previously reported regeneration-related genes (Supplementary Table 3), and then we investigated the expression patterns of these genes along the reconstructed redifferentiation trajectory. As shown in Fig. 3a,b, we observed 3 distinct dynamic expression patterns. Genes in patterns 1 and 2 had gradually increased and decreased expression trend, respectively, while genes in pattern 3 had specifically high expression in cell populations 3 (Fig. 3a,b).

**Fig. 3.**
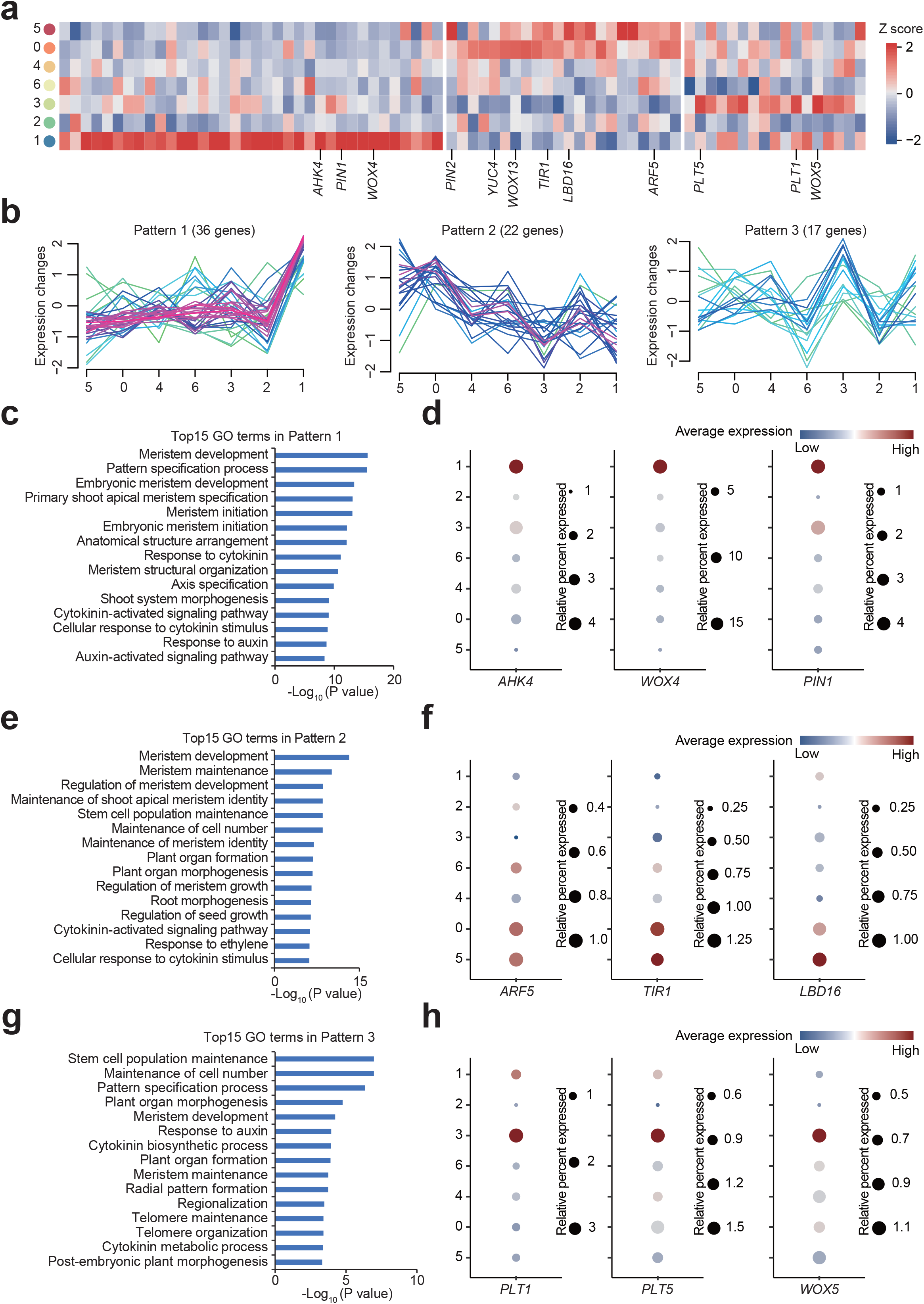
Dynamic expression patterns of canonical genes during *de novo* shoot regeneration. **a**, The heatmap of 75 canonical genes in *de novo* shoot organogenesis. Blue and red represent z score < 0 and > 0, respectively. **b**, Dynamic expression patterns of 75 canonical genes in *de novo* shoot organogenesis along the development trajectory. **c**, Top 15 GO terms in pattern 1. **d**, Dynamic expression of represented *AHK4, WOX4*, and *PIN1* genes in pattern 1. **e**, Top 15 GO terms in pattern 2. **f**, Dynamic expression of represented *ARF5, TIR1*, and *LBD16* genes in pattern 2. **g**, Top 15 GO terms in pattern 3. **h**, Dynamic expression of represented *PLT1/5* and *WOX5* genes in pattern 3. Relative Percent Expressed means the percentage of the number of gene expression cells in all cells of a specific cluster relative to the percentage of expression in cluster 5. The average expression means the average expression of genes at the single-cell level in specific clusters.

Thirty-nine genes were present in pattern 1, and these genes were highly expressed in cell population 1. GO analysis showed that the genes related to the primary shoot apical meristem specification, meristem initiation, and shoot system morphogenesis were highly enriched in pattern 1 (Fig. 3c), and the finding was consistent with the fact that the regenerating callus would eventually form the shoot. Among the 39 genes in pattern 1, some genes have been confirmed as key factors for shoot pro-meristem formation, such as cytokinin receptor genes *AHK4, WOX4*, and *PIN1* ^44,45^ (Fig. 3d).

For pattern 2, 22 genes were present, and they were highly expressed on day 0 (Fig. 3b). We found that the genes related to biosynthesis, polar transportation, receptor, and response factor of auxin were included in pattern 2, such as *ARF5, TIR1, LBD16, YUC4*, and *PIN2* (Fig. 3f), which was consistent with the enrichment of auxin biosynthetic process in cell populations 5 and 0 in Fig. 2e,f and Supplementary Fig. 5a. In addition, *ARF5, LBD16*, and *TIR1* are known as key factors in regulating callus formation^46,47^. We also detected the expression of *WOX13* in Pattern 2 (Supplementary Fig. 5a), and this gene has been verified to modify cell walls for efficient callus formation^48^. Considering many genes in pattern 2 have been reported to be involved in regulating callus formation, we speculated that these genes in pattern 2 could mainly play roles in regulating the formation of callus, but not in the shoot regeneration process^46,47^.

As for gene expression pattern 3, 17 genes were highly expressed in the identified QC-like cells, and the stem cell population maintenance GO term was especially enriched in this expression pattern (Fig. 3g). We found that genes involved in maintaining the quiescent central niche were included in this pattern, such as *PLT1/5*, and *WOX5* (Fig. 3h), indicating that these 17 genes might promote the acquisition of cell pluripotency during the redifferentiation process^49,50^. Taken together, here we demonstrate that the 75 canonical regeneration-related genes were specifically expressed at certain stages during shoot regeneration, and these findings shed light on the shoot regeneration mechanisms.

### Inference of cell type-specific transcription factor regulatory networks

To understand the roles of transcription factors (TFs) during the early stage of *de novo* shoot regeneration, we analyzed the expression of TFs in our dataset. Among the detected 1254 TFs we screened 101 TFs for further analysis (Supplementary Table 4), which were highly expressed in more than 33% cells of each cell population.

First, we investigated the expression patterns of the 101 transcription factors along the re-constructed redifferentiation trajectories. As shown in Fig. 4a, we observed three main expression patterns. For example, some TFs were highly expressed in cell populations 5 and 0, while other TFs were highly expressed in cell populations 3 and 1, respectively (Fig, 4a), indicating that different developmental stages require different sets of TFs during *de novo* shoot regeneration. Consistent with the above-mentioned results in this study (Fig. 3a and Supplementary Fig. 4b), we observed the high similarity between cell populations 5 and 0. Then we checked which TF was present in these patterns. We found that TFs such as *RAP2*.*6, NAP, ATMYBR1*, and *HY5* were highly expressed in cell populations 5 and 0, and *WRKY70, WRKY18*, and *WRKY75* were predominantly expressed in cell population 3, and *WRKY18* and *RRTF1* were highly expressed in the cell population 1. As far as we know, these TFs have not been proved to be involved in the regeneration process. Based on the dynamic and specific expression patterns, these TFs could play important roles in regulating the cell fate transition during *de novo* shoot regeneration, and further mutant analysis is required in future studies

**Fig. 4.**
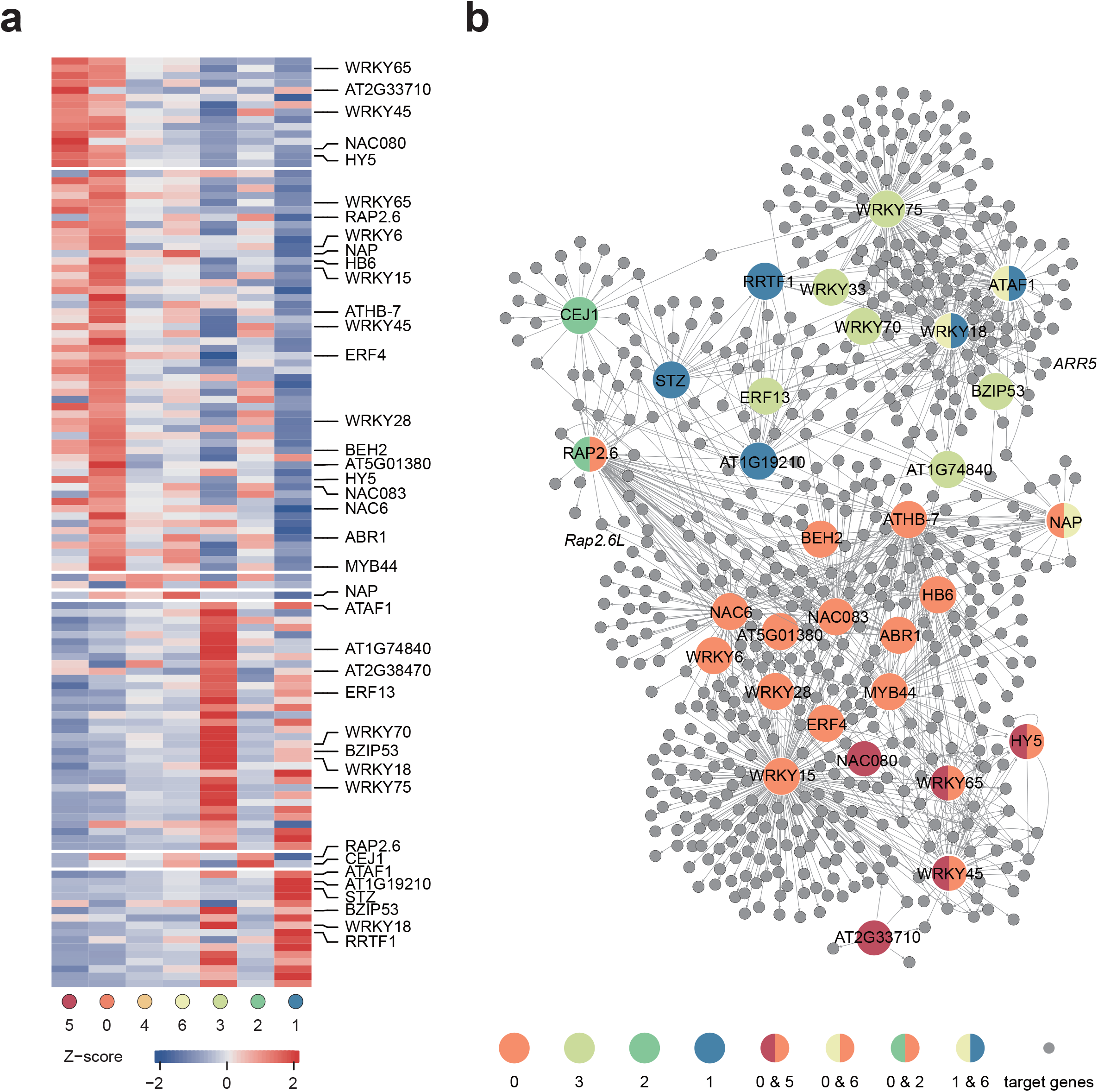
Transcription factor (TF) regulatory network analysis at the early stage of *de novo* shoot organogenesis. **a**, Heatmap of upregulated marker TFs for each cluster. **b**, The regulatory networks of 31 core TFs. Marker TFs in clusters 5, 0, 4, 6, 3, 2, and 1 were represented by red, orange, yellow, canary yellow, grass green, green, and blue circles, respectively.

To further explore the TFs and their target genes, we performed TF regulatory network (TFRN) analysis. By combining the regulatory relationships analysis results and the co-expression analysis with the verified *Arabidopsis* cistrome^51^, we were able to show the TFRNs (Fig. 4b). 31 core TFs and 642 target genes were revealed (Supplementary Table 4). Five and 17 TFRNs were identified in cell populations 5 and 0, respectively, and 3 TFRNs were shared between these two cell populations, such as *WRKY45, WRKY65*, and *HY5*. For the QC-like cell population 3, 8 TFRNs were identified, and *ATAF1, ERF13, bZIP53, WRKY33, WRKY70, WRKY18, WRKY75*, and *AT1G74840* were included. Five TFRNs were identified in cell population 1, such as *ATAF1, ZAT10, WRKY18, RRTF1*, and *AT1G19210*. Among these TFRNs, *RAP2*.*6L* is required for *de novo* shoot regeneration^52,53^, and we discovered that there were 3 candidate TFs that directly regulate *RAP2*.*6L* expression, including *RAP2*.*6, MYB44*, and *VNI2*. In addition, *ARR5*, a transcription repressor that mediates the negative feedback loop in cytokinin signaling, showed a significantly increased expression pattern when callus was transferred to SIM (Supplementary Fig. 5b), which was consistent with the role of *ARR5* in pluripotency acquisition^10^. We found that the *WRKY18* might directly promote the expression of *ARR5* during *de novo* shoot regeneration. The predicted TFRNs provided valuable resources for future studies on the mechanism of *de novo* shoot regeneration.

## Discussion

Here using scRNA-seq, we characterized the early stage of *de novo* shoot regeneration from a root-derived callus and reconstructed the redifferentiation trajectories. The characteristics of the seven major cell populations along the redifferentiation trajectories would provide valuable resources for a comprehensive and deeper understanding of the mechanism of shoot organogenesis.

To construct the developmental trajectory of callus cells during *de novo* shoot regeneration, we innovatively selected pluripotency-related genes as input for cell clustering^54^. We successfully constructed the redifferentiation trajectory which corresponded to the biological developmental process. Although we tried to apply tools, such as monocle 2 and 3, for constructing the developmental trajectory, we did not obtain reasonable results (Supplementary Fig. 2a-e), indicating that suitable tools or methods need to be selected for different datasets.

Along the developmental trajectory, we found that cells had relatively higher pluripotency at the starting points compared with cells at a later developmental stage. Interestingly, we identified the QC-like cells at the intermediate stage of redifferentiation, and the considerably increased expression of well-known QC regulator, *WOX5*, in this cell cluster further supported the QC-like cell identity. Consistently, QC-like cells, which have been suggested to be primarily responsible for organ regeneration^10^, have also been identified during lateral root formation^21^. QC cells are the core cells of Root apical meristem (RAM) and could generate diverse root cell types^55^. Correspondingly, we detected the pericycle-like cells appeared at the later stage. In addition, cell cluster 1 also showed similarity to several other root cell types, indicating that the redifferentiation directions might diverge at this point and further studies are needed to uncover the complex developmental processes. Moreover, we found that the JA and SA-related biological processes were enriched and several TFs were highly expressed in the QC-like cells, which provided valuable insights for further research to reveal the roles of QC-like cells during *de novo* shoot regeneration.

In addition to phytohormones, light is another important factor for regulating *de novo* shoot regeneration. In this study, the adding of light treatment was an essential factor for promoting shoot regeneration after callus was transferred to SIM. Here we found that *HY5*, a basic leucine zipper (bZIP) transcription factor and a positive regulator of photomorphogenesis, was highly expressed at the beginning of redifferentiation trajectories, which was consistent with the finding that the loss of function in *HY5* leads to the defect of callus formation^56^. However, the expression of *HY5* was gradually decreased in the following stages, indicating that *HY5* might play its role during the formation of callus. In addition to *HY5*, we also found other light-related genes, including *miR163* and its target *PXMT1*, which might also be involved in the *de novo* shoot regeneration process. Previously, it has been reported that light-inducible *miR163* targets *PXMT1* transcripts to regulate *Arabidopsis* seed germination and primary root elongation^57^. In this study, when the callus was transferred to SIM and cultured on the light condition, *miR163* was expressed immediately, while its target *PXMT1* that highly expressed at the previous stage and could not be detected (Supplementary Fig. 5c). The inversely correlated expression pattern of these two genes indicated that the expression of *miR163* is induced by light and it further targets *PXMT1* transcripts to regulate the process of *de novo* shoot regeneration. Collectively, we speculate that light-inducible *miR163* and its targets *PXMT1* are involved in regulating the *de novo* shoot regeneration process, and further verifications are needed.

The reconstructed redifferentiation trajectories, the dynamic gene expression, and the complex regulatory networks at single-cell resolution provided great insights and valuable resources for future studies on *de novo* shoot regeneration. In this study, we only revealed the early stage of *de novo* shoot regeneration. To have an overall view of *de novo* shoot regeneration, single-cell transcriptome profiling of cells after day 10 is still needed. Considering the friable callus gradually changes into compact tissues, the collection of single cells by protoplasting is no longer infeasible. Therefore, using single-nucleus RNA-seq should be a better solution. Moreover, epigenetic regulations play important roles in regulating the regeneration process, and the single-cell sequencing assay for transposase-accessible chromatin (scATAC-seq) is the state-of-the-art technology for analyzing genome-wide chromatin accessibility at the single-cell level during *de novo* shoot regeneration.

## Methods

### Plant growth conditions and *de novo* shoot regeneration culturing

*A. thaliana* Col-0 was used as the wild-type. Seeds were sterilized with (0.05% (v/v) Tween-20 solution for 10 minutes) and subsequently sown on 1/2 Murashige and Skoog (MS) medium (Sigma M5519) at 22□ in long-day conditions (16 h light/8 h dark). Harvested root explants from 5-days (d)-old seedlings were cultured on callus-induction medium (CIM) with MS medium, 3% w/v sucrose, 0.3%w/v Phytagel, 0.22 μM N-(Phenylmethyl)-9H-purin-6-amine and 4.52 μM 2,4-Dichlorophenoxyacetic acid, buffered to PH 5.7, at 25□ in dark conditions for callus induction, and sub-culture was performed every two weeks. Two months later, calli were collected and transferred to shoot induction medium (SIM) with MS medium, 1% w/v sucrose, 0.05% w/v MES (Sigma, M3671), 0.3% w/v Phytagel, 0.1% v/v Gambor’s B5 vitamin, 9.12 μM zeatin, 4.1 μM biotin and 1.97 μM 3-indolebutyric acid, buffered to PH 5.7, at 22□ in 16 h light/8 h dark conditions for shoot regeneration.

### Protoplasting

Protoplast isolation was performed with a modified version of the polyethylene glycol (PEG)-mediated method as described previously^58^. Samples were collected from two-month-old CIM-induced calli in 0, 2, 4, 6, and 10 days after being transferred into SIM medium. All the calli were digested in the protoplast isolation solution containing 1% w/v Cellulase ‘Onozuka’ R–10 (Yakult Pharmaceutical), 0.2%w/v macerozyme R-10 (Yakult Pharmaceutical), 0.1% w/v Pectinase (Sigma, 17389), 20 mM MES (pH 5.7, Sigma, M3671), 10 mM CaCl_2_ (Sigma, C5670), 10 mM KCl (Sigma, P5405), 0.4 M D-mannitol (Sigma, M1902), and 0.1% BSA (Sigma, V900933) for approximately 2 h on a shaker at 60 rpm in the dark respectively. To remove enzyme solution and tissue debris, the digested tissue suspensions were filtered through 40 μm nylon mesh and washed by W5 buffer with the components of 154 mM NaCl (Sigma, S5886), 5 mM KCl (Sigma, P5405), 125 mM CaCl_2_, and 2 mM MES (pH 5.7). The mixed solutions were gently centrifugated at 100 g for 5 min to collect Protoplasts, which were then washed and resuspended with W5 buffer at a concentration of 2 × 10^6^ cells/mL, counted using a hemocytometer.

### scRNA-seq

Single-cell RNA-seq libraries were prepared with a DNB C4 scRNA-seq kit following the published protocol^59,60^. In brief, 100 ul W5 buffer containing protoplasts was supplemented with 6% ficoll 400, 0.04% BSA, 2U/ul RNase inhibitor as single-cell suspension of every sample, which was loaded into the chip with Barcoded mRNA capture beads and oil for droplet generation. After incubating at room temperature for 20 minutes, collected droplets were broken by the bead filter and the bead pellets were resuspended with 100 μl RT mix in the thermal cycle as follows: 42 °C for 90 minutes, 10 cycles of 50 °C for 2 minutes, 42 °C for 2 minutes. The bead pellets were then resuspended in 200 μl of exonuclease mix for digestion and amplified by PCR as follows: 95 °C for 3 27 minutes, 15 cycles of 98 °C for 20 s, 58 °C for 20 s, 72 °C for 3 28 minutes, and finally 72 °C for 5 minutes. Amplified cDNA was purified and subsequently fragmented for indexed sequencing libraries according to the manufacturer’s protocol of the DNB C4 scRNA-seq kit. All libraries were further prepared based on the BGISEQ-T1 sequencing platform. The DNA nanoballs (DNBs) were loaded into the patterned nanoarrays and sequenced on the BGISEQ-T1 sequencer using the following read length: 41 bp for read 1, 100 bp for read 2, and 10 bp for sample index.

### Bulk RNA-seq data processing

Total RNA was exacted from two-month-old CIM-induced calli in 0, 2, 4, and 6 days after being transferred into SIM medium using miRNeasy Mini Kit (QIAGEN, 217004). High-quality RNA was used to prepare sequencing libraries with the MGIEasy RNA library preparation kit. All libraries were sequenced on the BGISEQ-T1 sequencer using the following read length: 100 bp for read 1, 100 bp for read 2, and 10 bp for sample index. Then, analysis of the bulk RNA-seq data was performed. First, the bulk RNA-seq data were filtered using SOAPnuke (v1.5.2) with parameters -l 5 -q 0.2 -n 0.05^61^. After removing low-quality bases RNAs, the clean data were mapped to the *Arabidopsis thaliana* genome (TAIR10) by using HISAT2^62^ with parameters “-k 1 -p 4 -x --min-intronlen 40 --max-intronlen 5000 --no-unal --dta --un-gz --rna-strandness RF”. Then the expression levels of each gene were calculated by the transcripts per kilobase of exon model per million mapped reads (TPM) by using StringTie^63^ with parameters “-t -C -e -B -A --rf”. The final TPM matrix of all samples was used for subsequent analysis.

### Single-cell RNA-seq data processing

First, single-cell data were mapped to *Arabidopsis thaliana* genome (TAIR10) by using STAR^64^ with parameters “--outStd SAM --outSAMunmapped Within --runThreadN 4 --alignIntronMin 40 --alignIntronMax 5000”. Then, the gene expression level in each cell is calculated by the unique molecular identifiers (UMI) using PISA (https://github.com/pisa-engine/pisa).

In addition, we also downloaded the 10x single-cell RNA-seq data of *Arabidopsis thaliana* root (GSM4466788 and GSM4466787 of GSE122687)^27^ and shoot (CRS098237)^29^ from GEO and GSA, respectively. First, fastq-dump was used to convert SRA data of root into fastq format. Then these 10X sequencing data were mapped to the *Arabidopsis thaliana* genome (TAIR10) and generated the UMI matrix of each cell by using Cell Ranger (version 5.0.1) count pipeline.

### Identification of genes related to pluripotency and differentiation

Seurat’s “findMarkers” function was used to identify the differentially expressed genes (DEGs) between root and shoot and D0, respectively. These differential genes were defined as root-up, D0d-up1, shoot-up, and D0d-up2, respectively. The overlapped genes of D0d-up1 and D0d-up2 were defined as pre-pluripotency-related genes. And the shoot-up genes were defined as pre-differentiation-related genes. At the same time, some key genes in the regeneration process were collected, and these genes were defined as key genes. The Mfuzz^65^ function was also used to find gene sets that showed the same trend from Day 0 to Day 6/10 in bulk and single-cell data from the three gene sets. On Day 0 to Day 6/10, the gene set with a decreasing trend was defined as pluripotency-related genes, while the gene set with an increasing trend was defined as differentiation-related genes. Finally, the four genes were combined to obtain the genes related to callus development.

### Clustering

The cells with less than 500 genes, more than 5,000 genes, and more than 10 mitochondria were filtered out. Through cell quality control, 35,669 high-quality cells were finally obtained for subsequent analysis. The genes related to callus development were selected for linear dimensionality reduction using the function “RunPCA”. The “FindNeighbors” and “FindClusters” functions were used for cell clustering respectively, with parameters of 30 PC and 0.2 resolution. Finally, the “RunUMAP” function was used for visualization.

### Gene signature scores

First, each Gene in a given gene set was normalized by the “scale” function in all cells. Then, the normalized scores of gene set in each cell were averaged, which was defined as Gene signature scores.

### Differential expression analysis

The cell identity scores were calculated using Hypergeometric Distribution Enrichment Analysis (HDEA). Specifically, the root cell type-specific markers from the online database^38^ and the leaf cell type-specific markers from Arabidopsis shoot apex^29^ were used as reference (Supplementary Table 2). The cluster-specific markers were calculated using Seurat “FindAllMarkers” function log-fold change >= 0.25, present in >= 25% cells. The z-score of genes from specific GO terms was calculated.

### GO enrichment analysis

GO enrichment analysis was performed using R package clusterProfiler^66^ with TAIR10 annotation as the background. The smaller the P-value is, the more the GO term is significantly enriched. The GO terms enriched in the top 100 markers (using all markers if markers were less than 100) for each cluster were performed.

### Correlation analysis

The “cor” function was used to perform correlation analysis on the whole transcriptome of each single cell sample. The distance used was Euclidian, and the cluster method was “kendall” from the normalized data for each sample calculated by Seurat.

### Transcription factor (TF) regulatory network analysis

Firstly, genes with more than 33% gene expression in the marker list of each cluster were defined as candidate genes, and then the transcription factor list of each cluster was screened from these genes. The GENIE3^67^ was used to predict the regulatory relationships of these transcription factors and candidate genes from the normalized data for each cell calculated by Seurat with parameters “nTrees = 1000, nCores = 8” and the random seed was 123. Then the top 10,000 regulatory relationships were screened for subsequent analysis. And the known regulatory relationships between transcription factors (TFs) and target genes of *Arabidopsis* were downloaded from the Plant Cistrome Database^51^. Finally, the regulatory relationships present in both the database and the top 10,000 regulatory relationships predicted by GENIE3^67^ were obtained.

### Trajectory analysis

Trajectory analysis was performed using R packages Monocle2 and Monocle3^37^, respectively. Among them, the key genes used in Monocle2 were the top 770 highly variable genes. The default parameters were used for analysis in Monocle2 and Monocle3.

### Statistical analysis

Statistical analysis was performed using R (version 4.0.4). Graphs were generated using R package (ggplot2, mfuzz, ComplexHeatmap, and cytoscape). The Wilcoxon rank-sum test was used to calculate differential genes between different cell types or samples, and p < 0.05 was considered significant. Fisher’s Exact Test was used for gene function annotation.

## Acknowledgments

This research was supported by Guangdong Provincial Key Laboratory of Genome Read and Write (No. 2017B030301011). We also sincerely thank the support provided by China National Gene Bank (CNGB).

## Author contributions

K.X. and H.-X.S. designed and supervised the study. K.X. and G.L., designed the experiment, K.X., G.L., Z.C., P.Y., R.Y., and X.L. performed the protoplast isolation, library preparation, and sequencing. H.-X.S., J.L., J.-M.L., and R.C. performed bioinformatics analysis. Z.C., P.Y., and Z.L. performed the tissue culture experiment. G.L., J.-M.L., and J.L. wrote the manuscript. K.X., H.-X.S., and Y.G. participated in the manuscript editing and discussion. All authors edited and approved the manuscript.

## Competing interests

The authors declare that there is no conflict of interests.

## Data availability

The sequencing data of Arabidopsis thaliana that support the findings of this study have been deposited into CNGB Sequence Archive (CNSA)^68^ of China National GeneBank DataBase (CNGBdb)^69^ with accession number CNP0002343.

## Supplementary Information

**Supplementary Fig. 1** | **The data quality analysis of callus at the early stage of *de novo* shoot regeneration. a**, The 2-month-old root-derived calli grew on SIM for 26 days. **b**, Correlation analysis of 15 scRNA-seq samples. **c**, Gene number, UMI, percentage of mitochondrial and chloroplast genes at the single-cell level at five time points. **d**, Correlation analysis of gene expression between bulk RNA-seq and scRNA-seq data in Day 0, 2, 4, and 6.

**Supplementary Fig. 2** | **Developmental trajectory analysis using Monocle2 and Monocle3. a**-**b**, Development trajectory using Monocle2. **c**, Developmental trajectory split by time points using Monocle2. **d**-**e**, Development trajectory using Monocle3.

**Supplementary Fig. 3** | **Selecting genes for reconstructing the developmental trajectory. a**, Relatively increased expression pattern (p) of the pre-pluripotency-related genes in the bulk (b) RNA-seq and scRNA-seq (sc) data in Day 0, 2, 4, 6 and 10. **b**, Relatively decreased expression patterns (p) of the pre-pluripotency-related genes in the bulk (b) RNA-seq and scRNA-seq (sc) data in Day 0, 2, 4, 6 and 10. **c**, Genes with both decreased expression patterns in both bulk RNA-seq and scRNA-seq were designated as pluripotency-related genes, while genes with both increased expression pattern in both bulk RNA-seq and scRNA-seq were designated as differentiation-related genes. **d**, Regeneration-related genes with decreased expression patterns in both bulk RNA-seq and scRNA-seq were selected. **e**, Regeneration-related genes with relatively increased expression patterns in both bulk RNA-seq and scRNA-seq were chosen.

**Supplementary Fig. 4** | **Callus cell type characterization. a**, Expression of phase-specific genes at single-cell level during *de novo* shoot regeneration, including *PLT1, PLT2*, and *ESR1* marker genes. **b**, Heatmap of top 100 GO terms in 7 cell clusters. **c**, The specific hormone-related GO term in cluster 0 and 5.

**Supplementary Fig. 5** | **Dynamic expression patterns of specific genes. a**, Dynamic expression of *WOX13, YUC4*, and *Pin2* in pattern 2 of Fig. 3b. **b**, Dynamic expression pattern of *ARR5*. **c**, Dynamic expression pattern of *PXMT1* and *ath-MIR163*. Relative Percent Expressed means the percentage of the number of gene expression cells in all cells of a specific cluster relative to the percentage of expression in cluster 5. The average expression means the average expression of genes at the single-cell level in specific clusters.

## Notes

### Competing Interest Statement

The authors have declared no competing interest.

## References

1 Radhakrishnan, D. et al. Shoot regeneration: a journey from acquisition of competence to completion. Curr Opin Plant Biol 41, 23–31, doi:10.1016/j.pbi.2017.08.001 (2018).

2 Efferth, T. Biotechnology applications of plant callus cultures. Engineering 5, 50–59, doi:10.1016/j.eng.2018.11.006 (2019).

3 Maher, M. F. et al. Plant gene editing through de novo induction of meristems. Nat Biotechnol 38, 84–89, doi:10.1038/s41587-019-0337-2 (2020).

4 Debernardi, J. M. et al. A GRF–GIF chimeric protein improves the regeneration efficiency of transgenic plants. Nature Biotechnology, doi:10.1038/s41587-020-0703-0 (2020).

5 Shin, J., Bae, S. & Seo, P. J. De novo shoot organogenesis during plant regeneration. J Exp Bot 71, 63–72, doi:10.1093/jxb/erz395 (2020).

6 Zhang, T.-Q. et al. A two-step model for de novo activation of WUSCHEL during plant shoot regeneration. The Plant Cell 29, 1073–1087, doi:10.1105/tpc.16.00863 (2017).

7 Meng, W. J. et al. Type-B ARABIDOPSIS RESPONSE REGULATORs specify the shoot stem cell niche by dual regulation of WUSCHEL. The Plant Cell 29, 1357–1372, doi:10.1105/tpc.16.00640 (2017).

8 Liu, J. et al. The WOX11–LBD16 pathway promotes pluripotency acquisition in callus cells during de novo shoot regeneration in tissue culture. Plant and Cell Physiology 59, 739–748, doi:10.1093/pcp/pcy010 (2018).

9 Kareem, A. et al. PLETHORA genes control regeneration by a two-step mechanism. Current Biology 25, 1017–1030, doi:10.1016/j.cub.2015.02.022 (2015).

10 Zhai, N. & Xu, L. Pluripotency acquisition in the middle cell layer of callus is required for organ regeneration. Nature Plants 7, 1453–1460, doi:10.1038/s41477-021-01015-8 (2021).

11 Aida, M., Ishida, T. & Tasaka, M. Shoot apical meristem and cotyledon formation during Arabidopsis embryogenesis: interaction among the CUP-SHAPED COTYLEDON and SHOOT MERISTEMLESS genes. Development 126, 1563–1570, doi:10.1242/dev.126.8.1563 (1999).

12 Matsuo, N., Makino, M. & Banno, H. Arabidopsis ENHANCER OF SHOOT REGENERATION (ESR) 1 and ESR2 regulate in vitro shoot regeneration and their expressions are differentially regulated. Plant science 181, 39–46, doi:10.1016/j.plantsci.2011.03.007 (2011).

13 Banno, H., Ikeda, Y., Niu, Q.-W. & Chua, N.-H. Overexpression of Arabidopsis ESR1 induces initiation of shoot regeneration. The Plant Cell 13, 2609–2618, doi:10.1105/tpc.010234 (2001).

14 Cheng, Z. J. et al. Pattern of Auxin and Cytokinin Responses for Shoot Meristem Induction Results from the Regulation of Cytokinin Biosynthesis by AUXIN RESPONSE FACTOR3 Plant Physiology 161, 240–251, doi:10.1104/pp.112.203166 (2013).

15 Ikeuchi, M., Ogawa, Y., Iwase, A. & Sugimoto, K. Plant regeneration: cellular origins and molecular mechanisms. Development 143, 1442, doi:10.1242/dev.134668 (2016).

16 Chupeau, M. C. et al. Characterization of the early events leading to totipotency in an Arabidopsis protoplast liquid culture by temporal transcript profiling. Plant Cell 25, 2444–2463, doi:10.1105/tpc.113.109538 (2013).

17 Mironova, V. & Xu, J. A single-cell view of tissue regeneration in plants. Current Opinion in Plant Biology 52, 149–154, doi:https://doi.org/10.1016/j.pbi.2019.09.003 (2019).

18 Zheng, H.-x., Wu, F.-h., Li, S.-m., Zhang, X. S. & Sui, N. Single-cell profiling lights different cell trajectories in plants. aBIOTECH 2, 64–78, doi:10.1007/s42994-021-00040-7 (2021).

19 Efroni, I. et al. Root Regeneration Triggers an Embryo-like Sequence Guided by Hormonal Interactions. Cell 165, 1721–1733, doi:https://doi.org/10.1016/j.cell.2016.04.046 (2016).

20 Zhang, T.-Q., Chen, Y., Liu, Y., Lin, W.-H. & Wang, J.-W. Single-cell transcriptome atlas and chromatin accessibility landscape reveal differentiation trajectories in the rice root. Nature Communications 12, 2053, doi:10.1038/s41467-021-22352-4 (2021).

21 Serrano-Ron, L. et al. RECONSTRUCTION OF LATERAL ROOT FORMATION THROUGH SINGLE-CELL RNA-SEQ REVEALS ORDER OF TISSUE INITIATION. Molecular Plant, doi:10.1016/j.molp.2021.05.028 (2021).

22 Gala, H. P. et al. A single cell view of the transcriptome during lateral root initiation in Arabidopsis thaliana. The Plant Cell, doi:10.1093/plcell/koab101 (2021).

23 Denyer, T. et al. Spatiotemporal developmental trajectories in the Arabidopsis root revealed using high-throughput single-cell RNA sequencing. Developmental cell 48, 840–852. e845 (2019).

24 Jean-Baptiste, K. et al. Dynamics of gene expression in single root cells of Arabidopsis thaliana. The Plant Cell 31, 993–1011 (2019).

25 Liu, Q. et al. Transcriptional landscape of rice roots at the single-cell resolution. Molecular Plant 14, 384–394 (2021).

26 Ryu, K. H., Huang, L., Kang, H. M. & Schiefelbein, J. Single-cell RNA sequencing resolves molecular relationships among individual plant cells. Plant physiology 179, 1444–1456 (2019).

27 Shulse, C. N. et al. High-throughput single-cell transcriptome profiling of plant cell types. Cell reports 27, 2241–2247. e2244 (2019).

28 Zhang, T.-Q., Xu, Z.-G., Shang, G.-D. & Wang, J.-W. A single-cell RNA sequencing profiles the developmental landscape of Arabidopsis root. Molecular plant 12, 648–660 (2019).

29 Zhang, T.-Q., Chen, Y. & Wang, J.-W. A single-cell analysis of the Arabidopsis vegetative shoot apex. Developmental Cell 56, 1056–1074.e1058, doi:10.1016/j.devcel.2021.02.021 (2021).

30 Satterlee, J. W., Strable, J. & Scanlon, M. J. Plant stem-cell organization and differentiation at single-cell resolution. Proceedings of the National Academy of Sciences 117, 33689–33699, doi:10.1073/pnas.2018788117 (2020).

31 Bezrutczyk, M. et al. Evidence for phloem loading via the abaxial bundle sheath cells in maize leaves. The Plant Cell 33, 531–547 (2021).

32 Kim, J.-Y. et al. Distinct identities of leaf phloem cells revealed by single cell transcriptomics. The Plant Cell 33, 511–530 (2021).

33 Lopez-Anido, C. B. et al. Single-cell resolution of lineage trajectories in the Arabidopsis stomatal lineage and developing leaf. Developmental Cell 56, 1043–1055.e1044, doi:10.1016/j.devcel.2021.03.014 (2021).

34 Liu, Z. et al. Global Dynamic Molecular Profiles of Stomatal Lineage Cell Development by Single-Cell RNA Sequencing. Molecular Plant 13, 1178–1193, doi:10.1016/j.molp.2020.06.010 (2020).

35 Yu, W., Qh, A., Ke, L. A. & Wqa, B. Single-cell transcriptome atlas of the leaf and root of rice seedlings. Journal of Genetics and Genomics, doi:10.1016/j.jgg.2021.06.001 (2021).

36 Wendrich, J. R. et al. Vascular transcription factors guide plant epidermal responses to limiting phosphate conditions. Science 370, eaay4970, doi:10.1126/science.aay4970 (2020).

37 Qiu, X. et al. Single-cell mRNA quantification and differential analysis with Census. Nature methods 14, 309–315, doi:10.1038/nmeth.4150 (2017).

38 Chen, H. et al. PlantscRNAdb: A database for plant single-cell RNA analysis. Molecular Plant, doi:10.1016/j.molp.2021.05.002 (2021).

39 Sugimoto, K., Jiao, Y. & Meyerowitz, E. M. Arabidopsis regeneration from multiple tissues occurs via a root development pathway. Developmental cell 18, 463–471, doi:10.1016/j.devcel.2010.02.004 (2010).

40 Lavenus, J. et al. Lateral root development in Arabidopsis: fifty shades of auxin. Trends Plant Sci 18, 450–458, doi:10.1016/j.tplants.2013.04.006 (2013).

41 Skoog, F. & Miller, C. O. Chemical regulation of growth and organ formation in plant tissues cultured in vitro. Cheminform 27, 118, doi:DOI: 10.1146/annurev-arplant-050718-100434 (1996).

42 Achard, P. et al. The plant stress hormone ethylene controls floral transition via DELLA-dependent regulation of floral meristem-identity genes. Proceedings of the National Academy of Sciences 104, 6484–6489, doi:10.1073/pnas.0610717104 (2007).

43 Su, Y. H., Tang, L. P., Zhao, X. Y. & Zhang, X. S. Plant cell totipotency: Insights into cellular reprogramming. Journal of Integrative Plant Biology 63, 228–243, doi:https://doi.org/10.1111/jipb.12972 (2021).

44 Kubiasová, K. et al. Cytokinin fluoroprobe reveals multiple sites of cytokinin perception at plasma membrane and endoplasmic reticulum. Nature communications 11, 1–11, doi:10.1038/s41467-020-17949-0 (2020).

45 Hao, Z. et al. Conserved, divergent and heterochronic gene expression during Brachypodium and Arabidopsis embryo development. Plant Reproduction, 1–18, doi:10.1007/s00497-021-00413-4 (2021).

46 Smit, M. E. et al. Specification and regulation of vascular tissue identity in the Arabidopsis embryo. Development 147, dev186130, doi:10.1242/dev.186130 (2020).

47 Goh, T. et al. Lateral root initiation requires the sequential induction of transcription factors LBD16 and PUCHI in Arabidopsis thaliana. New Phytologist 224, 749–760, doi:10.1242/dev.186130 (2019).

48 Momoko, I. et al. Wound-inducible WUSCHEL-RELATED HOMEOBOX 13 is required for callus growth and organ reconnection. Plant Physiology, doi:10.1093/plphys/kiab510 (2021).

49 Wang, Z. et al. Salicylic acid promotes quiescent center cell division through ROS accumulation and down-regulation of PLT1, PLT2, and WOX5. J Integr Plant Biol 63, 583–596, doi:10.1111/jipb.13020 (2021).

50 Wein, A., Le Gac, A.-L. & Laux, T. Stem cell ageing of the root apical meristem of Arabidopsis thaliana. Mechanisms of ageing and development 190, 111313, doi:10.1016/j.mad.2020.111313 (2020).

51 Niwa, H. The principles that govern transcription factor network functions in stem cells. Development 145, dev157420, doi:10.1242/dev.157420 (2018).

52 Yang, S., Poretska, O. & Sieberer, T. ALTERED MERISTEM PROGRAM1 restricts shoot meristem proliferation and regeneration by limiting HD-ZIP III-mediated expression of RAP2. 6L. Plant physiology 177, 1580–1594, doi:10.1104/pp.18.00252 (2018).

53 Che, P., Lall, S., Nettleton, D. & Howell, S. H. Gene expression programs during shoot, root, and callus development in Arabidopsis tissue culture. Plant physiology 141, 620–637, doi:10.1104/pp.106.081240 (2006).

54 Schiebinger, G. et al. Optimal-transport analysis of single-cell gene expression identifies developmental trajectories in reprogramming. Cell 176, 928–943. e922, doi:10.1016/j.cell.2019.01.006 (2019).

55 Drisch, R. C. & Stahl, Y. Function and regulation of transcription factors involved in root apical meristem and stem cell maintenance. Frontiers in Plant Science 6, 505, doi:10.3389/fpls.2015.00505 (2015).

56 Cluis, C. P., Mouchel, C. F. & Hardtke, C. S. The Arabidopsis transcription factor HY5 integrates light and hormone signaling pathways. The Plant Journal 38, 332–347, doi:10.1111/j.1365-313X.2004.02052.x (2004).

57 Chung, P. J. et al. Light-inducible miR163 targets PXMT1 transcripts to promote seed germination and primary root elongation in Arabidopsis. Plant physiology 170, 1772–1782, doi:10.1104/pp.15.01188 (2016).

58 Yoo, S. D., Cho, Y. H. & Sheen, J. Arabidopsis mesophyll protoplasts: a versatile cell system for transient gene expression analysis. Nature protocols 2, 1565–1572, doi:10.1038/nprot.2007.199 (2007).

59 Zhu, L. et al. Single-cell sequencing of peripheral mononuclear cells reveals distinct immune response landscapes of COVID-19 and influenza patients. Immunity 53, 685–696. e683, doi:10.1016/j.immuni.2020.07.009 (2020).

60 Ding, R. et al. Single-cell transcriptome analysis of the heterogeneous effects of differential expression of tumor PD-L1 on responding TCR-T cells. Theranostics 11, 4957, doi:10.7150/thno.55075 (2021).

61 Wang, Z., Gerstein, M. & Snyder, M. RNA-Seq: a revolutionary tool for transcriptomics. Nature reviews genetics 10, 57–63, doi:10.1038/nrg2484 (2009).

62 Kim, D., Langmead, B. & Salzberg, S. L. HISAT: a fast spliced aligner with low memory requirements. Nature methods 12, 357–360, doi:10.1038/nmeth.3317 (2015).

63 Pertea, M. et al. StringTie enables improved reconstruction of a transcriptome from RNA-seq reads. Nature Biotechnology 33, 290–295, doi:10.1038/nbt.3122 (2015).

64 Magoč, T. & Salzberg, S. L. FLASH: fast length adjustment of short reads to improve genome assemblies. Bioinformatics 27, 2957–2963, doi:10.1093/bioinformatics/btr507 (2011).

65 Futschik, M. E. & Carlisle, B. Noise-robust soft clustering of gene expression time-course data. Journal of bioinformatics and computational biology 3, 965–988, doi:0.1142/S0219720005001375 (2005).

66 Love, M. I., Huber, W. & Anders, S. Moderated estimation of fold change and dispersion for RNA-seq data with DESeq2. Genome biology 15, 1–21, doi:0.1186/s13059-014-0550-8 (2014).

67 Huynh-Thu, V. A., Irrthum, A., Wehenkel, L. & Geurts, P. Inferring regulatory networks from expression data using tree-based methods. PloS one 5, e12776, doi:10.1371/journal.pone.0012776 (2010).

68 Guo, X. et al. CNSA: a data repository for archiving omics data. Database 2020, doi:10.1093/database/baaa055 (2020).

69 Chen, F. Z. et al. CNGBdb: China National GeneBank DataBase. Yi chuan = Hereditas 42, 799–809, doi:10.16288/j.yczz.20-080 (2020).

