## Supplementary figures and images for "Single-cell transcriptome reveals the redifferentiation trajectories of the early stage of *de novo* shoot regeneration in *Arabidopsis thaliana*"

### Supplementary Fig. 1

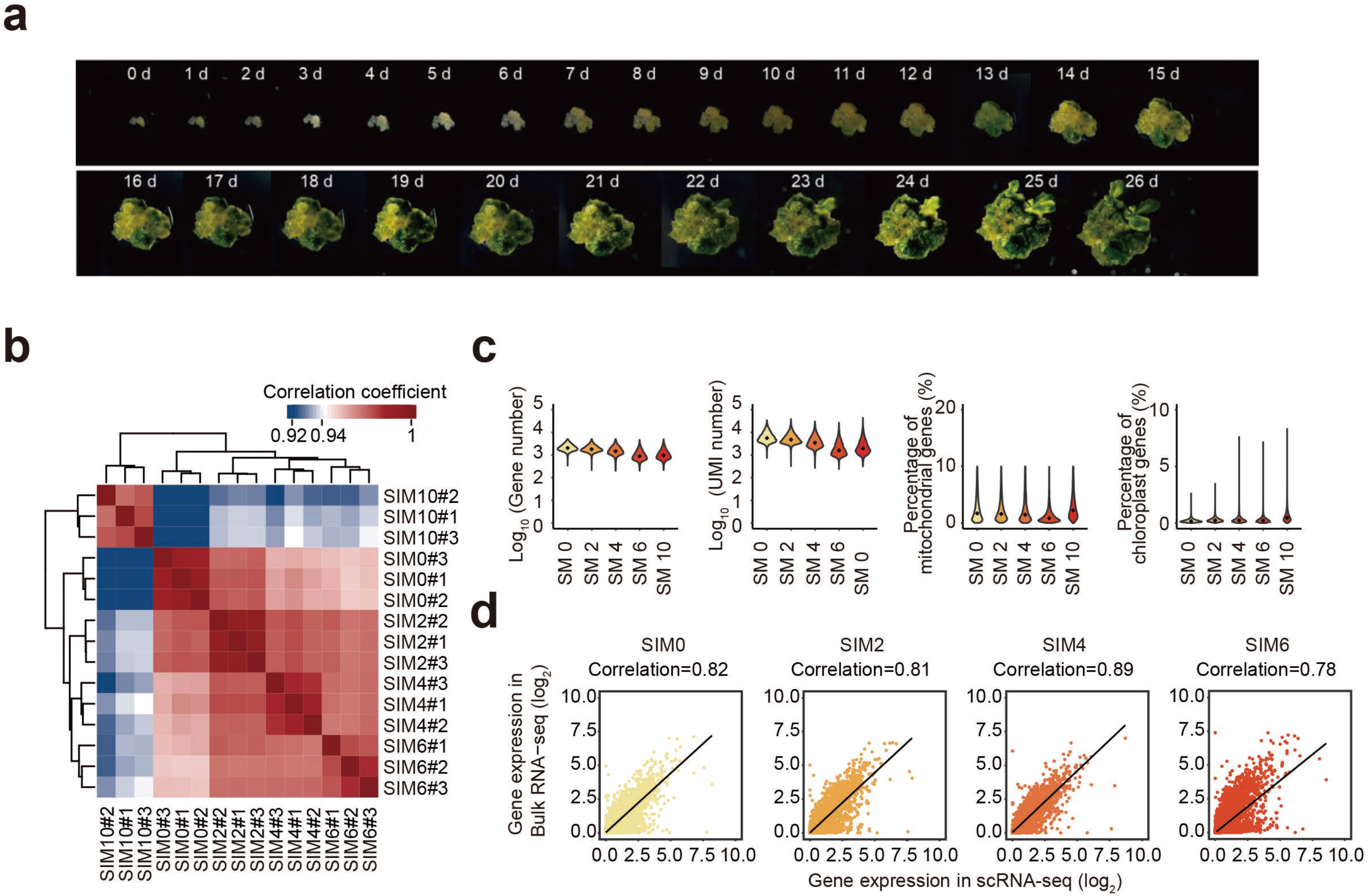

### Supplementary Fig. 2

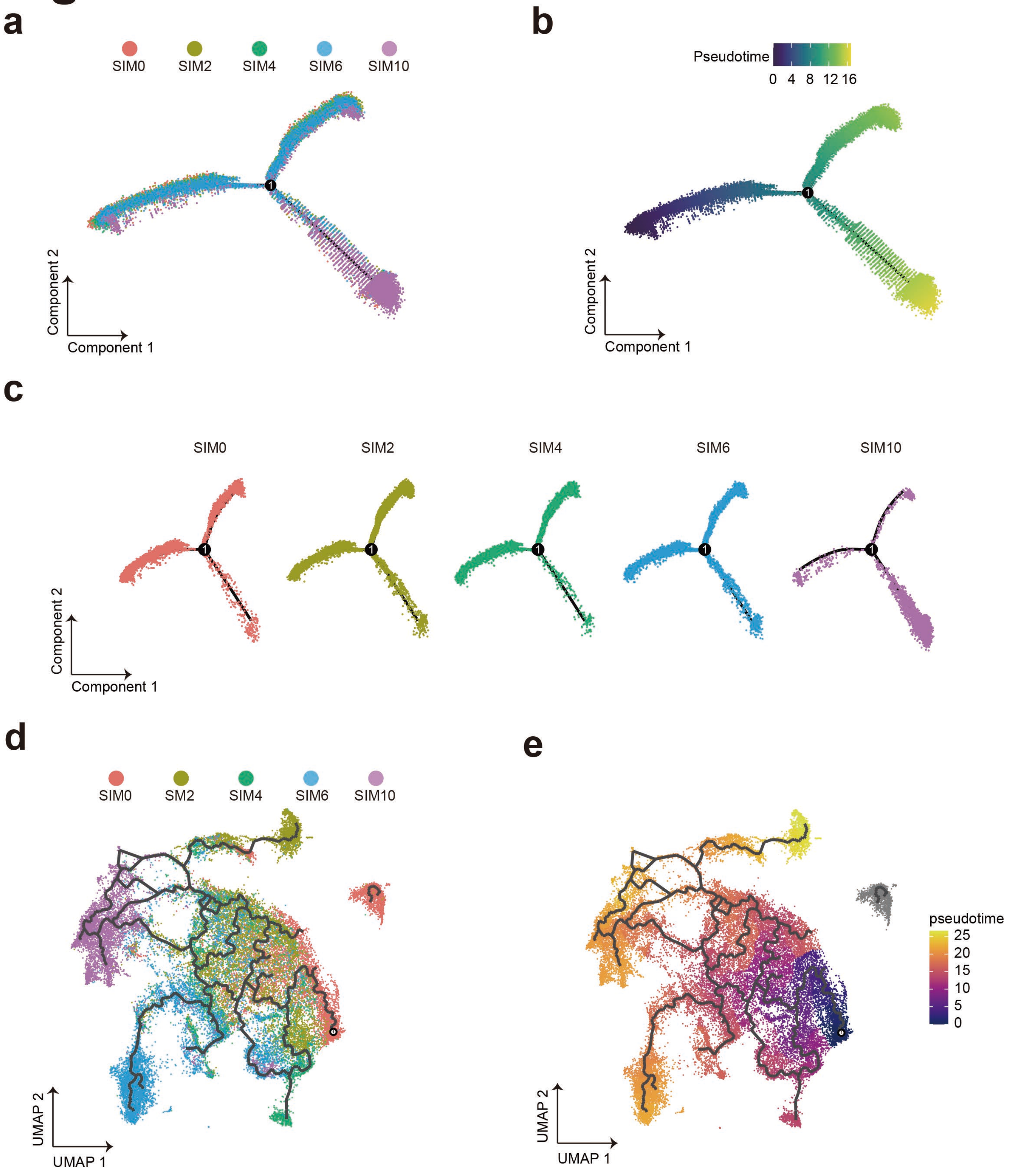

### Supplementary Fig. 3

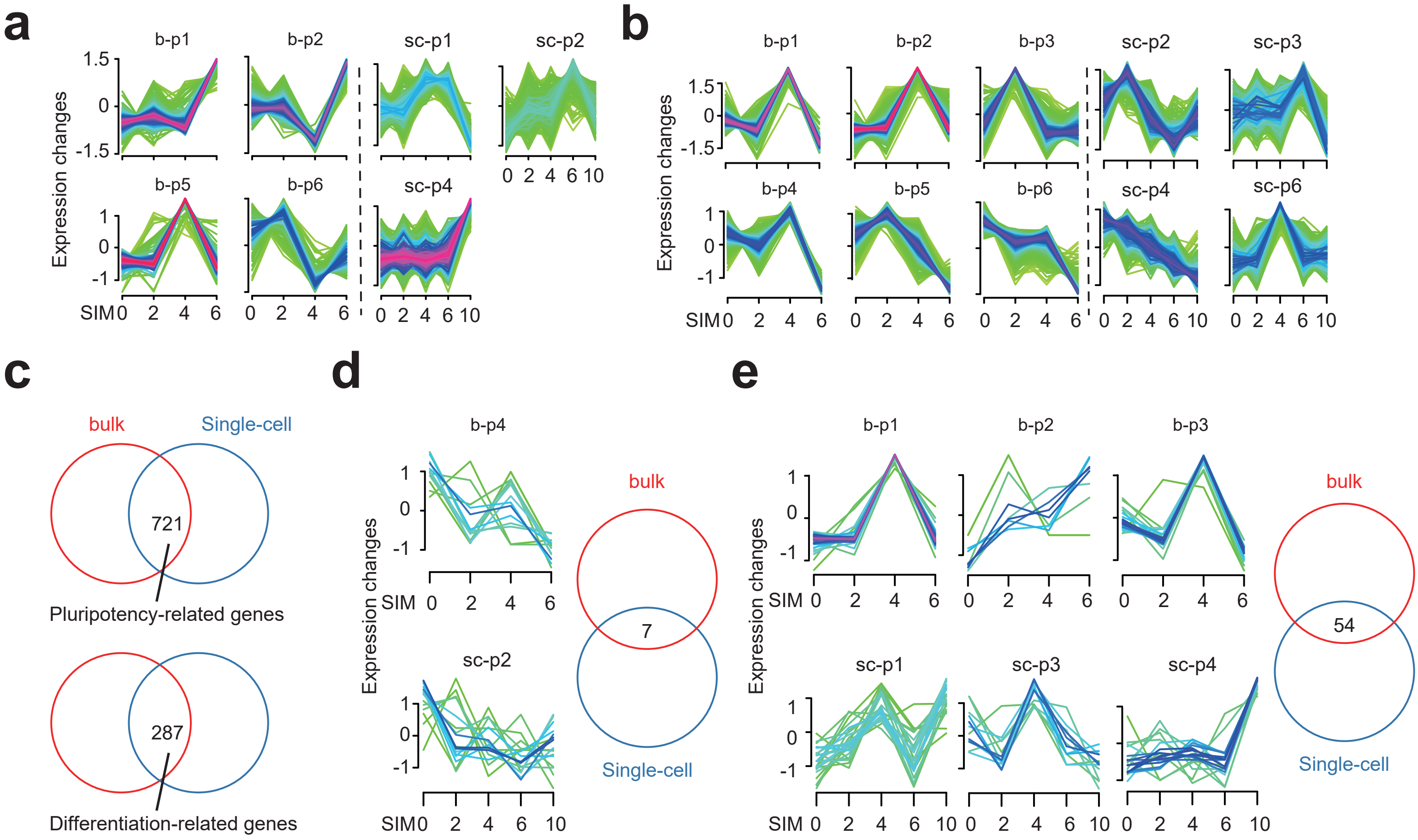

### Supplementary Fig. 4

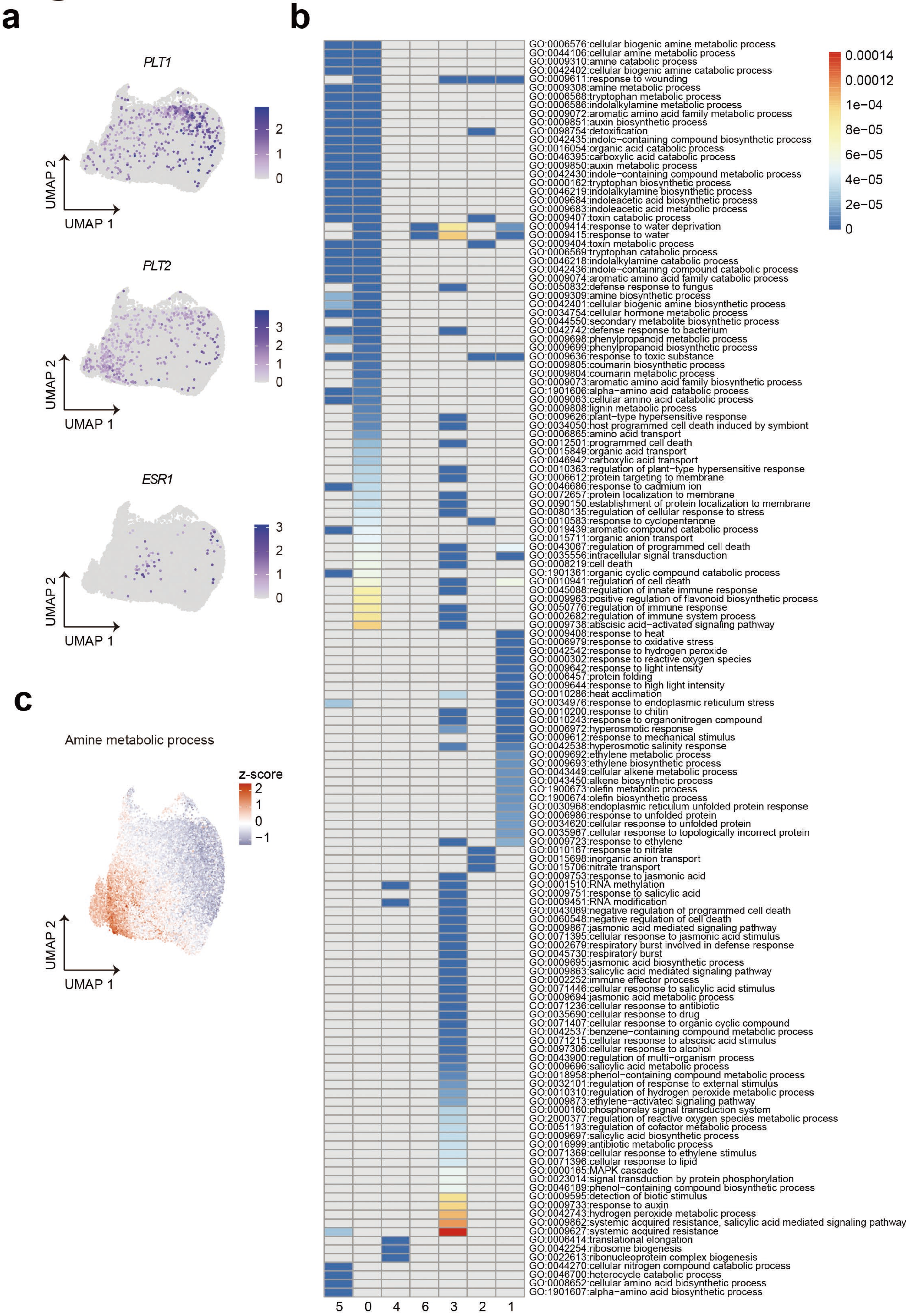

### Supplementary Fig. 5

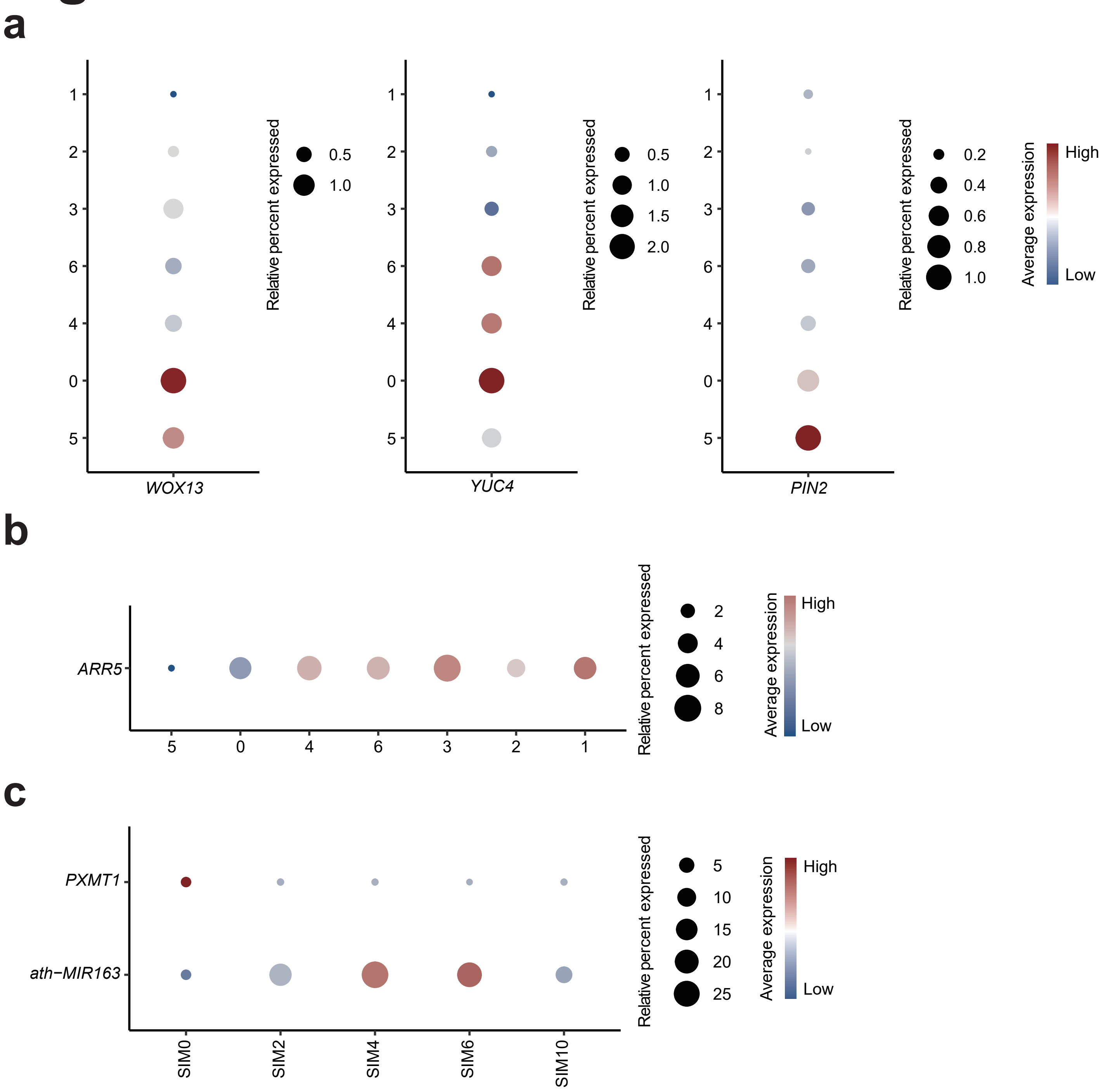
